# Natural variation in stomata size contributes to the local adaptation of water-use efficiency in *Arabidopsis thaliana*

**DOI:** 10.1101/253021

**Authors:** H. Dittberner, A. Korte, T. Mettler-Altmann, A.P.M. Weber, G. Monroe, J. de Meaux

## Abstract

Stomata control gas exchanges between the plant and the atmosphere. How natural variation in stomata size and density contributes to resolve trade-offs between carbon uptake and water-loss in response to local climatic variation is not yet understood. We developed an automated confocal microscopy approach to characterize natural genetic variation in stomatal patterning in 330 fully-sequenced *Arabidopsis thaliana* accessions collected throughout the European range of the species. We compared this to variation in water-use efficiency, measured as carbon isotope discrimination (δ^13^C). We detect substantial genetic variation for stomata size and density segregating within *Arabidopsis thaliana*. A positive correlation between stomata size and δ^13^C further suggests that this variation has consequences on water-use efficiency. Genome-wide association analyses indicate a complex genetic architecture underlying not only variation in stomata patterning but also to its co-variation with carbon uptake parameters. Yet, we report two novel QTL affecting δ^13^C independently of stomata patterning. This suggests that, in *A. thaliana*, both morphological and physiological variants contribute to genetic variance in water-use efficiency. Patterns of regional differentiation and co-variation with climatic parameters indicate that natural selection has contributed to shape some of this variation, especially in Southern Sweden, where water availability is more limited in spring relative to summer. These conditions are expected to favor the evolution of drought avoidance mechanisms over drought escape strategies.

## Introduction

In plants, carbon uptake and water loss are intimately linked by a trade-off between growth and water conservation (Cowan, 1986; Cowan & Farquhar, 1977; Field, Merino, & Mooney, 1983). Stomata, the microscopic pores embedded in the epidermis of plant leaves, play a key role in the resolution of this trade-off. Their density, distribution and regulation control the rate of CO_2_ and water exchange (Raven, 2002). As a result, they impact the ratio of photosynthetic carbon assimilation to water loss *via* transpiration. This ratio defines water-use efficiency (WUE), a physiological parameter that directly determines plant productivity when the water supply is limited. Variation in density, distribution and regulation of stomata may thus have played a pivotal role in shaping the diversity of plant communities throughout the globe (Lambers, Chapin, & Pons, 1998; McDowell et al., 2008).

The density of stomata on the leaf surface is expected to correlate positively with the rate of gas exchanges between the leaf and the atmosphere, also called “conductance”. Models based on gas diffusion theory predict that small stomata in high density can best maximize conductance (Franks & Beerling, 2009). A positive relationship between stomata density and conductance has been reported in a majority of studies looking at natural variation between species (Anderson & Briske, 1990; Pearce, Millard, Bray, & Rood, 2006) as well as within species (Carlson, Adams, & Holsinger, 2016; Muchow & Sinclair, 1989; Reich, 1984). Yet, higher stomata density does not always translate into higher rates of gas exchanges: in a diversity panel of rice (Ohsumi, Kanemura, Homma, Horie, & Shiraiwa, 2007) or within several vegetable crop species (Bakker, 1991), for example, the relationship was not observed.

Molecular mutants in genes promoting stomata development show that reduced stomata density translates into decreased water loss and increased ability to survive after exposure to drought (Franks, W. Doheny-Adams, Britton-Harper, & Gray, 2015; Yoo et al., 2010). Yet, decreased stomata density does not necessarily associate with increased demands on WUE imposed by water limitation. In the *Mimulus guttatus* species complex, accessions from drier inland populations showed decreased stomatal density and increased WUE, compared to accessions collected in humid coastal populations (Wu, Lowry, Nutter, & Willis, 2010). By contrast, in 19 *Protea repens* populations measured in a common garden experiment, stomata density increased with decreasing summer rainfall at the source location (Carlson et al., 2016).

In fact, stomata density is not the only parameter modulating the balance between water loss and carbon uptake. Variation in stomata size also impacts the efficiency of stomata regulation (Raven, 2014). Stomata open and close in response to environmental and internal signals (Chater et al., 2011; Kinoshita et al., 2011). This ensures that plants do not desiccate when water evaporation is maximal and spares water when photosynthesis is not active (Daszkowska-Golec & Szarejko, 2013). The speed of stomata closure is higher in smaller stomata (Drake, Froend, & Franks, 2013; Raven, 2014). Stomatal responses are an order of magnitude slower than photosynthetic changes, so any increase in closure time lag may result in unnecessary water loss and reduce WUE (T. Lawson, Kramer, & Raines, 2012; Raven, 2014). However, it is often observed that decreases in stomata size occur at the expense of increased stomata density (reviewed in Hetherington & Woodward, 2003). This leads to a correlation that may at first be counter-intuitive: an increase in stomata density can result in improved WUE because of indirect effects on stomata size. In *Eucalyptus globulus*, however, plants from the drier sites had smaller stomata and higher WUE but no concomitant change in stomata density (Franks, Drake, & Beerling, 2009). This suggested that the developmental effect correlating stomata size and density may sometimes be alleviated. Altogether, these studies highlight interconnections between stomata size, stomata density and WUE that change across species or populations. How and whether variation in these traits and their connections support or constrain adaptive processes, however, is not clearly established.

Eco-evolutionary studies, e.g. the analysis of evolutionary forces shaping genetic variation in natural populations, can determine whether phenotypic variance has a significant impact on the ecology of species (Carroll, Hendry, Reznick, & Fox, 2007; Hendry, 2016). By drawing on the elaborate toolbox of population genetics and genomics, it is not only possible to determine the genetic architecture of any given trait but also to ask whether it is optimized by natural selection and to investigate the ecological determinants of selective forces at work (Hendry, 2016; Weinig, Ewers, & Welch, 2014). In this effort, the annual species *Arabidopsis thaliana*, which thrives as a pioneer species in disturbed habitats, has a privileged position (Gaut, 2012). Genome-wide patterns of nucleotide variation can be contrasted to phenotypic variation and both the genetic architecture and the adaptive history of the traits can be reconstructed (Atwell et al., 2010; Fournier-Level et al., 2011; Alonso-Blanco et al., 2016). Environmental variation has a documented impact on local adaptation in this species (Debieu et al., 2013; Hamilton, Okada, Korves, & Schmitt, 2015; Hancock et al., 2011; Kronholm, Picó, Alonso-Blanco, Goudet, & Meaux, 2012; Lasky et al., 2014; Postma & Ågren, 2016). In addition, natural variation in stomatal patterning is known to segregate among *A. thaliana* accessions (Delgado, Alonso-Blanco, Fenoll, & Mena, 2011). This species thus provides the ideal evolutionary context in which the adaptive contribution of variation in stomata patterning can be dissected.

Here, we developed an automated confocal microscopy approach that overcomes the technical limitations which have so far complicated the phenotyping of stomatal variation on larger samples. We characterized genetic variation in stomatal patterning in 330 fully-sequenced accessions, across a North-South transect of the European range. Additionally, we measured δ^13^C, a commonly used estimate of water-use efficiency (WUE) (Juenger et al., 2005; Martin & Thorstenson, 1988; McKay et al., 2008; Mojica et al., 2016), for all genotypes. Combined with public genomic and environmental resources, this dataset allows us to ask: i) how variable are natural *A. thaliana* accessions in stomata patterning? ii) does variation in stomata patterning influence the carbon-water trade-off? iii) what is the genetic architecture of traits describing stomata patterning? iv) is stomata patterning optimized by natural selection? By combining a genome-wide association approach with Q_ST_/F_ST_ analyses and associations with environmental parameters, we show that, in *A. thaliana*, variation in stomata patterning plays a role in local adaptation. Our results further indicate that natural variation in stomata size is one of the adaptive traits contributing to the optimization of WUE.

## Methods

### Plant material, plant genotypes and growth conditions

In total, 330 accessions, spanning a wide geographical range were selected from the 1001 collection of fully sequenced genotypes (Suppl. Table 1). Accessions were assigned to five groups based on their geographic origin and genetic clustering (Alonso-Blanco et al., 2016): Spain, Western Europe, Central Europe, Southern Sweden and Northern Sweden (Figure S1). In 20 cases, for which genetic information contradicted geographic information, we prioritized geographic information since we are focusing on local adaptation and expect that geography, as opposed to demographic history, reflects the scale at which local adaptation proceeds. To avoid oversampling, we randomly reduced the number of plants sampled at the same location to one for the analysis of heritability, regional differentiation (Q_ST_-F_ST_) and climatic correlation, resulting in 287 accessions.

The genome sequences of the 330 genotypes included in the analysis were downloaded from the 1001 genome database (Alonso-Blanco et al., 2016) on May 12th, 2017. Single nucleotide polymorphism (SNP) data was extracted using *vcftools* (Danecek et al., 2011). Genomic data was thinned to 1 SNP picked randomly in each 1000bp window to reduce computational load. In *A. thaliana*, linkage disequilibrium extends beyond 1kb (Nordborg et al., 2002). Thus, this data-size reduction should not impact statistics describing the geographical structure of genomic variation. Additionally, minimum minor allele frequency was set to 5% and sites exceeding 5% missing data were removed, resulting in 70,410 SNPs among all genotypes. SNP information was loaded into R using the *vcfR* package (Knaus, Grunwald, Anderson, Winter, & Kamvar, 2017). For genome-wide association studies the full, unthinned SNP dataset was used and missing SNPs were imputed using BEAGLE version 3.0 (Browning & Browning, 2009).

Seeds were stratified on wet paper for 6 days at 4°C in darkness. Plants were grown on soil in 5×5 cm paper pots in 3 replicates with one plant per pot. Genotypes were randomized within each of 3 blocks of 12 trays containing 8×4 pots. Plants were grown for 7 weeks in growth chambers (one per block) under the following conditions: 16 h light; 95 µmol s^−1^ mm^−2^ light intensity; 20 °C day- and 18 °C night-temperature. Plants were watered twice a week and trays shuffled and rotated every two to three days to account for variable conditions within the chambers.

### High throughput phenotyping

After 7 weeks, one fully-expanded, intact, adult leaf (one of the largest leaves developed after leaf 4) was selected from each plant for microscopic analysis. Stomata density and size as well as leaf size were measured using our high-throughput microscopy pipeline (for details, see Suppl. Document 1). Stomata density was also determined manually on a random set of 14 individuals and on a set of 32 independently-grown individuals. Automatic and manual measurements were strongly correlated (Pearson correlation coefficient r^2^=0.88, p<<0.01and r^2^=0.81, p<<0.01, for the 14 and 32 individuals Figures S2-3). The algorithm was conservative and tended to slightly under-estimate stomata numbers, resulting in a low false-positive rate. This ensured that stomata area was generally quantified on objects that corresponded to real stomata. Due to quality filters in our pipeline, the number of analyzed images differed between samples (Figure S4). We found a significant correlation between the number of images analyzed and stomata density (r=0.21, p<<0.01, Figure S5), but not stomata size (r=0.02, p>0.05). Thus, we included the number of images as a co-factor into all statistical models for stomata density (see below). Carbon isotope discrimination measurements (δ^13^C of whole rosettes were performed for all plants in block 1 (for details see Suppl. Document 1).

### Heritability estimates

Broad-sense heritability *H^2^*, the proportion of the observed phenotypic variance that is genetic, was estimated as:

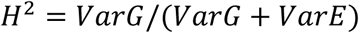

where VarG is the genetic variance and VarE is the environmental variance. Because we worked with inbred lines, VarE and VarG could be estimated as the variance between replicates of a genotype and the variance between genotypes, respectively, with a linear-mixed-model using block as fixed effect and genotype as random effect. We ran a linear mixed model using the *lme* function from *nlme* package (Pinheiro, Bates, DebRoy, Sarkar, & R Core Team, 2015) (Suppl. Document 2). δ^13^C, no replicates were available but a pseudo-heritability estimate was extracted from the GWAS mixed model including the kinship matrix (Atwell et al., 2010).

### Genome-Wide Association Study (GWAS)

For GWAS, SNPs with minor allele count <5 were removed, leaving a dataset of 2.8-3M SNPs, depending on missing data for the phenotypes. Minor allele frequency spectra for all three datasets show that the subset of 261 genotypes, for which all three phenotypes were determined, has a lower proportion of rare SNPs (Figure S6). GWAS was performed with a mixed model correcting for population structure using a kinship matrix calculated under the assumption of the infinitesimal model. SNPs were first analyzed with a fast approximation (Kang et al., 2010) and the 1000 top-most associated SNPs were reanalyzed with the complete model that estimates the respective variance components for each SNP separately (Kang et al., 2008).

For trait pairs measured on the same plant, a Multi-Trait Mixed Model (MTMM) was applied to distinguish common and trait-specific SNP-phenotype association (Korte et al., 2012). The MTMM performs three different statistical tests on a bivariate phenotype including each trait pair. The first model tests whether a given SNP has the same effect on both traits. This model has increased power to detect significant associations, which may fall under the significance threshold when traits are analyzed in isolation. The second model identifies SNPs having distinct effects on the two traits. It is well suited to detect SNPs with antagonistic effects on both traits. The last model combines both trait-specific and common effects. This last model is particularly powerful for detecting markers affecting both traits with different intensity. The MTMM analysis also provides estimates of the genetic and environmental correlation for each pair of traits. The statistical details of the models are described in (Korte et al., 2012).

For all analyses (GWAS and MTMM), the significance threshold for QTL identification was determined as a 5 % Bonferroni threshold, i.e. 0.05 divided by the number of SNPs in the dataset.

### Climatic data

Climatic data included average precipitation, temperature, water vapor pressure (humidity), wind speed and solar radiation estimates with 2.5 min grid resolution (WorldClim2 database (Fick & Hijmans, 2017) on May 30^th^, 2017) and soil water content (Trabucco & Zomer, 2010). For each variable and accession, we extracted a mean over the putative growing season, i.e. the months in the year with average temperature greater than 5 °C and average soil water content over 25% (Suppl. Table 1). We further computed historical drought frequencies at *A. thaliana* collection sites using 30+ years of the remotely-sensed Vegetative Health Index (VHI). The VHI is a drought detection method that combines the satellite measured Vegetative Health and Thermal Condition Indices to identify drought induced vegetative stress globally at weekly 4km^2^ resolution (Kogan, 1995). This is a validated method for detecting drought conditions in agriculture. Specifically, we used VHI records to calculate the historic frequency of observing drought conditions (VHI<40) during the spring (quarter surrounding spring equinox) and summer (quarter surrounding summer solstice). These are the typical reproductive seasons of *Arabidopsis* populations (reviewed in Burghardt, Metcalf, Wilczek, Schmitt, & Donohue, 2015). The drought regime in each location was quantified as the log-transformed ratio of spring over summer drought frequency. Positive values of this drought regime measure reflect environments where the frequency of drought decreases over the typical reproductive growing season, and vice versa for negative values. This ratio quantifies the seasonality of water availability. It correlates with the ratio of soil water content of the first and third month of the reproductive season (r=0.54, p<0.01), which we defined as the first and third growing month in the year, giving similar estimates as Burghardt, Metcalf, Wilczek, Schmitt, & Donohue (2015).

Because the seven climate variables are correlated, we combined them in seven principal components (PCs) for 316 *A. thaliana* collection sites (Figures S7-9, loadings described in Suppl. Document 2). Fourteen genotypes with missing climate data were excluded. Climatic distance between each region pair was estimated as the F-statistic of a multivariate analysis of variance (MANOVA) with climatic PCs as response variables and region of origin as predictor.

### Population genomic analysis

Principal component analysis (PCA) of genomic data (thinned to 1kb) was done using the *adegenet* package (Jombart et al., 2016) with missing data converted to the mean (Figures S10-11).

Comparing phenotypic differentiation (Q_ST_) to the distribution of F_ST_ is a useful method to reveal signatures of local adaptation (Leinonen, McCairns, O’hara, & Merilä, 2013; Whitlock & Guillaume, 2009). Genome-wide, pairwise F_ST_ estimates between regions were calculated using the *hierfstat* package (function *basic.stats*, Nei’s Fst) (Goudet, 2005). Negative F_ST_ values were set to zero before the 95th percentile was calculated.

For stomata density, stomata size and WUE, the respective phenotypic differentiation between regions, Q_ST_, was estimated as:

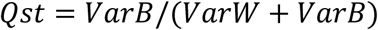

where VarW is the genetic variance within regions and VarB the genetic variance between regions as described in Kronholm et al. (2012) (for details, see Suppl. Document 1).

To test whether Q_ST_ estimates significantly exceed the 95^th^ percentile of the F_ST_ distribution, we permuted the phenotypic data by randomizing genotype labels to keep heritability constant. For each permutation and phenotype, we calculated the difference between each Q_ST_ value and the 95^th^ percentile of the F_ST_ distribution. We used the 95th percentile of the maximum Q_ST_-F_ST_ distance distribution as a threshold for determining if phenotypic differentiation significantly exceeds neutral expectations. Since this test takes the maximum Q_ST_-F_ST_ distance for all population combinations in each permutation, it does not require multiple testing correction.

### Statistical analysis

Statistical analysis was conducted using R (R Development Core Team, 2008) (R Markdown documentation in Suppl. Document 2). Plots were created using the following libraries: *ggplot2* (Wickham, 2009)*, ggthemes* (Arnold et al., 2017), *ggmap* (Kahle & Wickham, 2013)*, ggbiplot* (Vu, 2011) and *effects* (Fox et al., 2016).

We used Generalized Linear Models (GLM) to test the effect of block, origin, pot position in tray (edge or center) and leaf size on each phenotype (stomata density, stomata size and δ^13^C). For stomata density we also included the number of analyzed images as a co-factor. The error distribution was a quasi-Poisson distribution for stomata density and size and Gaussian for δ^13^C. Stomata density was log-transformed to avoid over-dispersion. Significance of each predictor was determined via a type-II likelihood-ratio test (*Anova* function of the *car* package). Significant differences between regions were based on GLMs including only significant predictor variables and determined with Tukey’s contrasts using the *glht* function of the *multcomp* package (Hothorn et al., 2017). GLMs were also used to test the impact of all climatic PCs on phenotypic traits, while accounting for population structure with the first 20 PCs for genetic variation, which explain 28% of genetic variation (see above). Additionally, for δ^13^C we also tested a simpler model including climatic parameters but not population structure. From the resulting models, we created effect plots for significant environmental PCs using the *effects* package (Fox et al., 2016). Further, we used GLMs with binomial distribution to test whether any of the climatic PCs significantly predicts the allelic states of loci associated with WUE in GWAS.

## Results

### Substantial genetic variation in stomata density and size

We analyzed over 31,000 images collected in leaves of 330 *A. thaliana* genotypes and observed high levels of genetic variation in stomata patterning. Genotypic means ranged from 87 to 204 stomata/mm^2^ for stomata density and from 95.0 µm^2^ to 135.1 µm^2^ for stomata size (see Suppl. Table 2 for raw phenotypic data). Leaf size was not significantly correlated with stomata density (r=-0.02, p=0.7, Figure S12) and stomata size (r=-0.08, p=0.15, Figure S13), as expected in fully developed leaves. Broad-sense heritability reached 0.41 and 0.30 for stomata size and density, respectively. Mean stomata density and stomata size were negatively correlated (r=-0.51, p<<0.001; Figure 1). Due to the strong correlation between stomata size and density, we focus primarily on stomata size in the following report, but results for stomata density are in the supplemental material.

**Figure 1:**
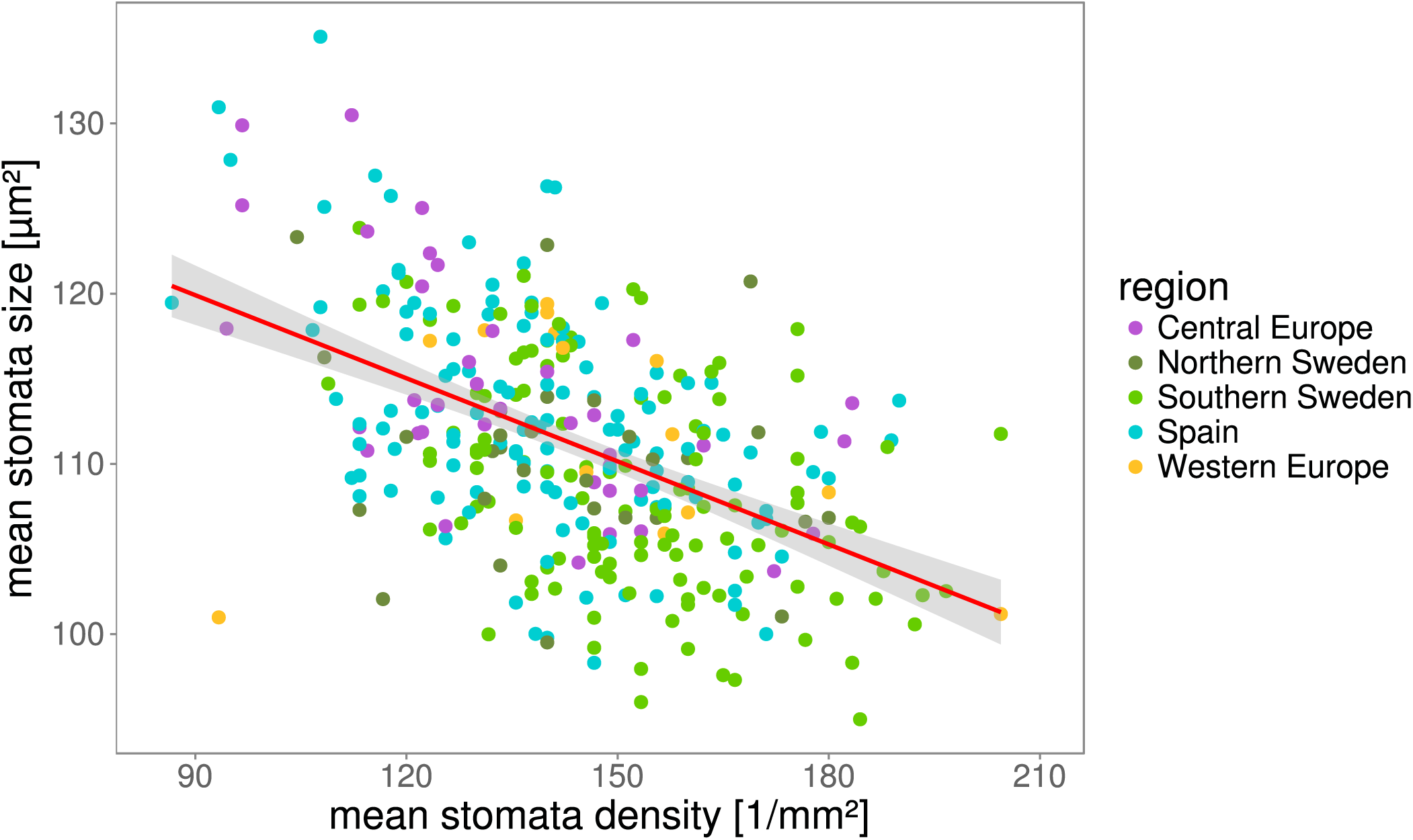
Natural variation in stomata patterning. Stomata density and size were measured for 330 natural genotypes of *A. thaliana*. The plot shows genotypic means of stomata density and stomata size. Dots are colored based on the geographical origin of each accession. The red line shows a linear fit and gray shadows indicate the error of the fit. Pearson’s product-moment correlation r=-0.5, p<0.001.

### Stomata size correlates with water-use efficiency

We expected variation in stomatal traits to influence the trade-off between carbon uptake and transpiration. Thus, we measured isotopic carbon discrimination, δ^13^C, an estimator that increases with water-use efficiency (WUE) (Farquhar, Hubick, Condon, & Richards, 1989; McKay et al., 2008). δ^13^C ranged from -38.7‰ to -30.8‰ (Suppl. Table 2) and was significantly correlated with stomata size (r=-0.18, p=0.004; Figure 2), indicating that accessions with smaller stomata have higher WUE. About ~4% of the total phenotypic variation (i.e. the sum of phenotypic and genetic variance) in δ^13^C is explained by genetic variance in stomata size. We found no significant correlation between stomatal density and δ^13^C (r=-0.007, p=0.9, Figure S14).

**Figure 2:**
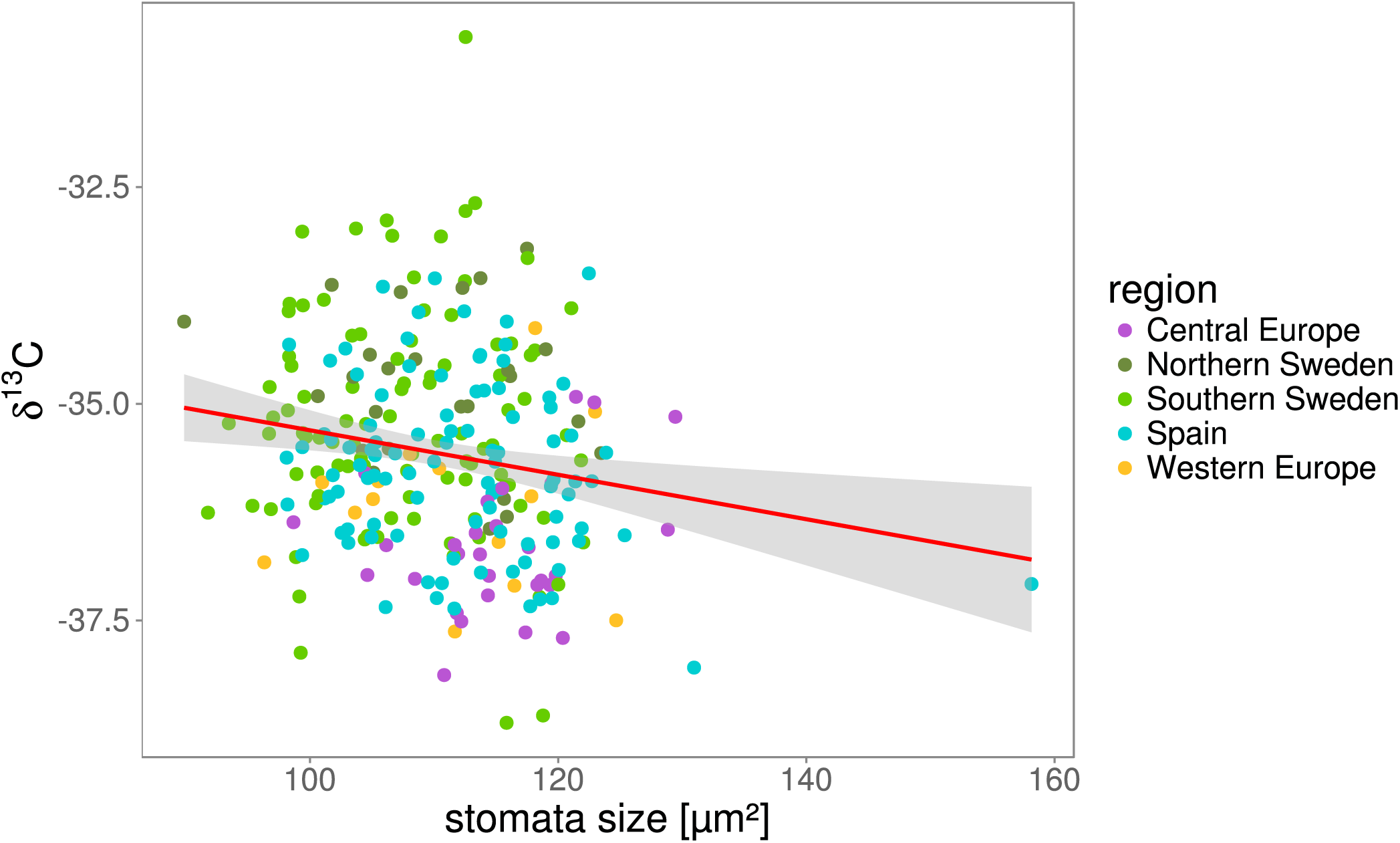
Stomata size correlates with water-use efficiency. δ^13^C was measured for all plants in block 1. Plots show correlation of stomata size (block 1 only) with δ^13^C. δ^13^C is expressed as ‰ against the Vienna Pee Dee Belemnite (VPDB) standard. The red line shows a linear fit and gray shadows indicate the error of the fit. Pearson’s product-moment correlation: r=-0.18, p=0.004. Correlation of δ^13^C and stomata size is not only driven by the Spanish outlier (correlation without outlier: r=-0.16, p=0.009). Genetic correlation was calculated using the MTMM approach: r=-0.58, p<0.05.

### Common genetic basis of stomata size and δ^13^C

To identify the genetic basis of the phenotypic variance we observe, we conducted a genome-wide association study (GWAS) for each phenotype. We calculated for each phenotype a pseudo-heritability, which is the fraction of phenotypic variance explained by the empirically estimated relatedness matrix (e.g. kinship matrix computed on genome-wide SNP typing). Pseudo-heritability estimates were 0.59 for stomata density, 0.56 for stomata size and 0.69 for δ^13^C, indicating that differences in stomata patterning and carbon physiology decreased with increasing relatedness. Despite considerable levels of heritability, we did not detect any variant associating with stomata density at a significance above the Bonferroni-corrected p-value of 0.05 (log10(p)=7.78). For stomata size, we detected one QTL with two SNPs significantly associating at positions 8567936 and 8568437 (Figure S15). These SNPs have an allele frequency of 1.5% (5 counts) and 2.1% (7 counts), respectively and map to gene *AT4G14990.1*, which encodes for a protein annotated with a function in cell differentiation. The former SNP is a synonymous coding mutation while the latter is in an intron.

For δ^13^C, one genomic region on chromosome 2 position 15094310 exceeded the Bonferroni significance threshold (log_10_(p)=7.97, Figure S16). Allele frequency at this SNP was 9.7% (30 counts) and all accessions carrying this allele, except four, were from Southern Sweden (3 Northern Sweden, 1 Central Europe). Southern Swedish lines carrying the allele showed significantly increased δ^13^C compared to the remaining Southern Swedish lines (W=1868, p-value=6.569e-05, Figure S17). A candidate causal mutation is a non-synonymous SNP at position 15109013 in gene *AT2G35970.1*, which codes for a protein belonging to the Late Embryogenesis Abundant (LEA) Hydroxyproline-Rich Glycoprotein family. This SNP also shows elevated association with the phenotype. However, its significance was below the Bonferroni-threshold (log(p)=7). Since this SNP is not in linkage disequilibrium with the highest associating SNP in the region (Figure S18), it is possible that another, independent SNP in this region is causing the association.

We used Multi-Trait Mixed-Model (MTMM) analysis to disentangle genetic and environmental determinants of the phenotypic correlations. We found that the significant correlation between stomata density and stomata size (r=-0.5) had no genetic basis, but had a significant (r=-0.9, p<0.05) residual correlation. This suggests that the correlation was not determined by common loci controlling the two traits, but by other, perhaps physical, constraints or by epistatic alleles at distinct loci. By contrast, the correlation between stomata size and δ^13^C (r= -0.18) had a significant genetic basis (kinship-based correlation, r=-0.58, p<0.05). Thus, in contrast to the phenotypic variation, genetic variation in stomata size roughly explains over 33% of the genetic variation in δ^13^C.

To further investigate the genetic basis for the correlation between stomata size and δ^13^C, we performed MTMM GWAS, which tests three models: the first model tests whether a SNP has the same effect on both traits; the second model tests whether a SNP has differing effects on both traits and the third model is a combination of the first two to identify SNPs which have effects of different magnitude on the traits (Korte et al., 2012). We did not observe variants with same or differing effects on δ^13^C and stomata size. However, with the combined model, we observed a marginally significant association on chromosome 4, which had an effect on δ^13^C but not stomata size. GWAS of δ^13^C restricted to the 261 individuals used for the MTMM analysis confirmed the QTL on chromosome 4. GWAS applied to different but overlapping sets of accessions yield similar results but can sometimes differ in the set of significant associations, since marginal changes in SNP frequency can affect significance levels (Figure S6). Indeed, the p-values of associations with δ^13^C for the two datasets (310 and 261 accessions) are highly correlated (r=0.87, p<<0.0001, Figure S19). In this set of genotypes, two SNPS, at position 7083610 and 7083612, exceeded the Bonferroni-corrected significance threshold (α=0.05) (both p=4.8e-09, Figure S20) although they were under the significance threshold in the larger dataset. Allele frequency is 14% (37 counts) at these two loci and explains 11% of the phenotypic variation. The association is probably due to complex haplotype differences since it coincides with a polymorphic deletion and contains several imputed SNPs. Thirty-five of the 37 accessions carrying the minor allele originated from Southern Sweden and showed significantly higher δ^13^C compared to other Southern Swedish accessions (mean difference=1.34; W=1707, p=1.15e-06; Figure S21). In summary, we detected two genetic variants significantly associating with δ^13^C, independent of stomata size, despite the common genetic basis of the two traits.

### Stomata size and stomata density correlate with geographical patterns of climatic variation

We used PCA to describe multivariate variation in climatic conditions reported for the locations of origins of the genotypes. We tested the correlation of each measured phenotype with climatic principal components (PCs) using a GLM which accounted for genetic population structure (see methods). We found a significant, negative relationship between genetic variation in stomata size and climatic PC2 (Likelihood ratio test Chi-Square (LRT X^2^)=9.2784, degrees of freedom (df)=1, p=0.005) and PC5 (LRT Χ^2^=5.7335, df=1, p=0.02, Figure 3). Climatic PC 2 explained 23.8% of climatic variation and had the strongest loadings (both negative) from temperature and water vapor pressure (humidity). Climatic PC 5 explained 9% of the climatic variation and mostly increased with increasing spring-summer drought probability ratio and increasing solar radiation. We also found significant climatic predictors for the distribution of genetic variation in stomata density (PC 2: LRT Χ^2^=8.6612, df=1 p=0.003; PC 5: LRT Χ^2^=7.3773, df=1, p=0.007; PC 7: LRT Χ^2^=6.6033, df=1, p=0.01; Figure S22). δ^13^C did not correlate with any of the climatic PCs. However, removing population structure covariates from the model revealed significant correlations of δ^13^C with climatic PC2 (+, LRT Χ^2^=7.3564, df=1, p=0.006), PC3 (-, LRT Χ^2^=3.8889, df=1, p=0.048) and PC4 (+, LRT Χ^2^=6.6885, df=1, p= 0.009) (Figure S23). PC3 explained 13.7% of climatic variation and principally increased with rainfall and decreased with spring-summer drought probability ratio. PC4 explained 11.4% of the total variation and mostly increased with wind speed. Therefore, the covariation of δ^13^C with climatic parameters describing variation in water availability and evaporation in *A. thaliana* is strong but confounded with the demographic history of the species. To test whether alleles associating with increased δ^13^C in GWAS are involved in adaptation to local climate, we checked whether any climatic PC is a significant predictor of the allelic state of Southern Swedish accessions. However, none of the climatic PCs was a significant predictor for one of the two loci.

**Figure 3:**
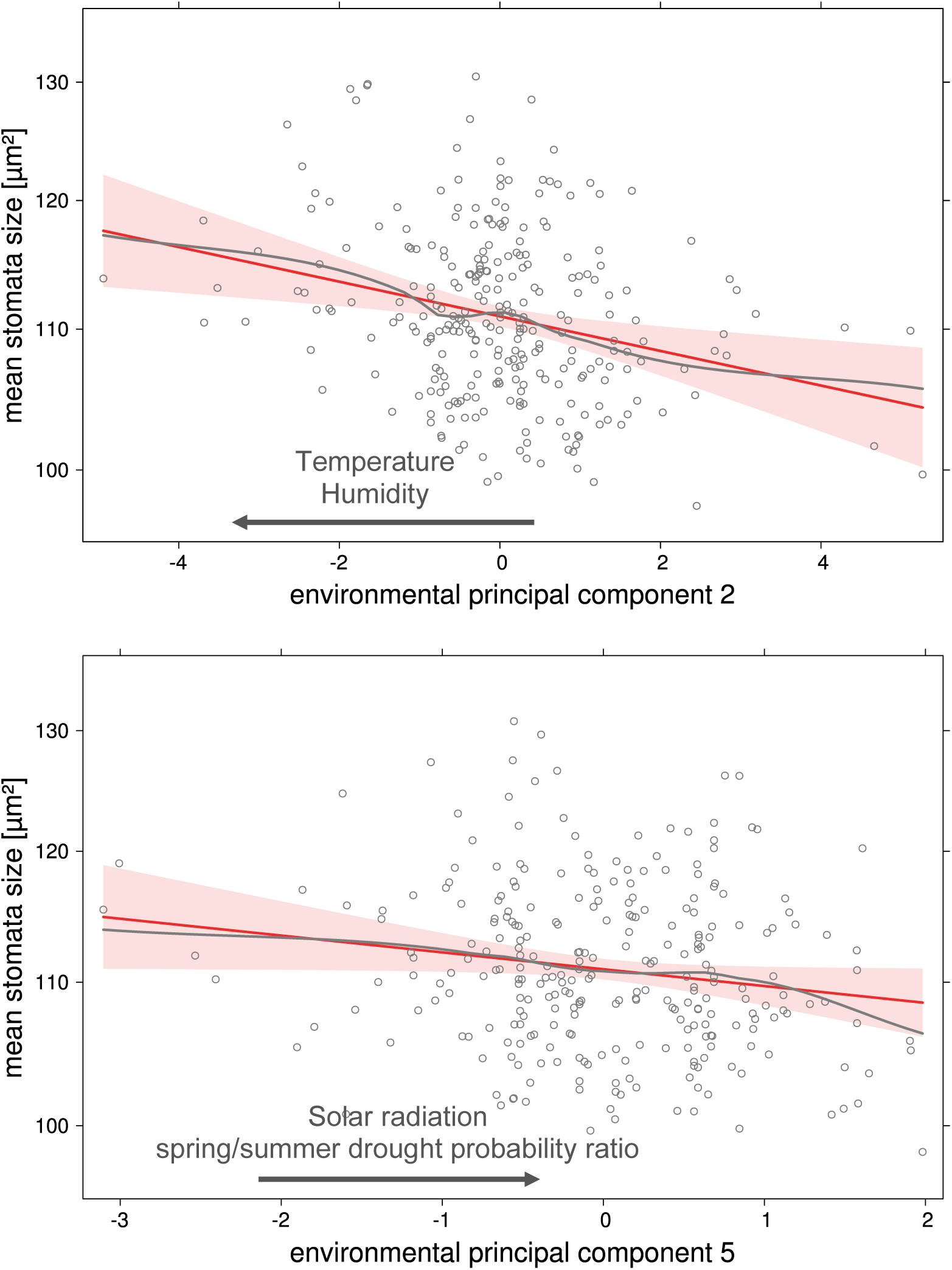
Stomata patterns correlate with geographical patterns of climatic variation. Correlation between stomata patterns and seven climatic principal components (PCs) was tested for each phenotype using a Generalized Linear Model (GLM) including genetic population structure as described by the 20 first genetic PCs. Plots are effect plots based on the GLM (see methods), showing the correlation between stomata size two climatic PCs. Black arrows indicate correlation with the climatic variables showing the strongest loadings for the respective PC. Plots show the linear fit (red solid line) and the smoothed fit of partial residuals (gray) of the specific predictor. Gray dots are partial residuals. The red shade shows the error of the linear fit. Both PCs shown here are significant predictors of the respective response variable (p<0.05).

### Patterns of regional differentiation depart from neutral expectations

Genotypes were divided into five regions based on genetic clustering (Alonso-Blanco et al., 2016) and their geographic origin (Figure S1, see Methods). We detected significant phenotypic differentiation among these regions for stomata size (LRT Χ^2^=52.852, df=4, p=9.151e-11, Figure 4). Stomata size was significantly lower in Southern Sweden (mean=108 µm^2^) compared to Central Europe (mean=114µm^2^, Generalized Linear Hypothesis Test (GLHT) z=-6.24, p<0.001), Western Europe (mean=111 µm^2^, GLHT z=2.769, p=0.04) and Spain (mean=113 µm^2^, GLHT z=6.709, p<0.001), which did not significantly differ from each other. Northern Sweden showed an intermediate phenotype and did not differ significantly from any region (mean=110 µm^2^). Variation for stomata density, showed a similar but inverted pattern (Figure S24).

**Figure 4:**
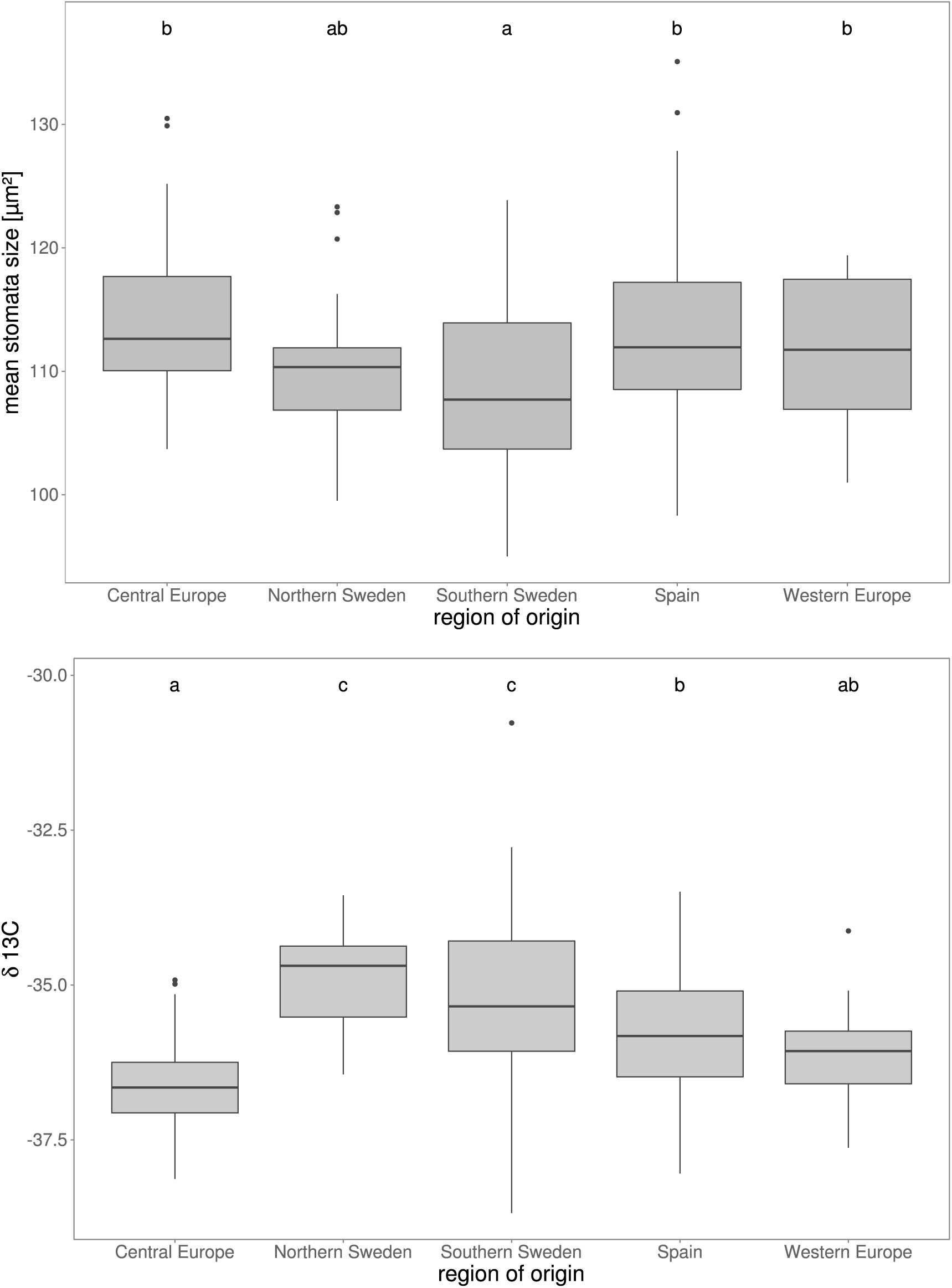
Significant regional differentiation of stomata size and δ^13^C. *A. thaliana* accessions were grouped based on their geographical origin. Boxplots show regional differentiation of stomata size (top) and δ^13^C (bottom). Significance of differentiation was tested using Generalized Linear Models followed by a post-hoc test. Statistical significance is indicated by letters on top: Groups that do not share a common letter are significantly different. Significance levels: top) a-c, a-bc: p<0.001; ab-c: p<0.05; bottom) a-c, a-b: p<0.001, b-c: p<0.01, ab-c: p<0.05.

Furthermore, we found significant regional differentiation in δ^13^C measurements (LR Χ^2^ =58.029, df=4 p=7.525e-12, Figure 4). Highest δ^13^C levels (highest WUE) were found in accessions from Northern Sweden (mean=-34.8) and Southern Sweden (mean=-35.2), which were significantly higher than in accessions from Spain (mean=-35.7; GLHT Southern Sweden z=-3.472, p=0.008; GLHT North Sweden z=-3.49, p=0.001) and Western Europe (mean=-36.06; GLHT Southern Sweden z= -2.8, p=0.03; GLHT Northern Sweden z=-3.28, p=0.008). Lowest δ^13^C levels were found in lines from Central Europe (mean=-36.6), which were significantly lower than in lines from Northern Sweden (GLHT z=5.676, p<0.001), Southern Sweden (GLHT z=6.992, p<0.001) and Spain (GLHT z=3.714, p=0.002).

The observed regional differences result either from the demographic history of the regions or from the action of local selective forces. To tease these possibilities apart, the phenotypic differentiation (Q_ST_) can be compared to nucleotide differentiation (F_ST_) (Kronholm et al., 2012; Leinonen et al., 2013). We examined each pair of regions separately, since they are not equidistant from each other and calculated F_ST_ distributions for over 70,000 SNP markers (spaced at least 1kb apart, see methods). For each trait, Q_ST_ exceeded the 95^th^ percentile of the F_ST_ distribution in at least two pairs of regions (Table 1 A-C). We used permutations to calculate a significance threshold for the Q_ST_/F_ST_ difference (see methods). Significant regional differentiation was pervasive in our sample, with Central Europe and Southern Sweden being significantly differentiated for all three phenotypes. This analysis suggests that natural selection has contributed to shape the phenotypic differentiation between regions.

**Table 1.**
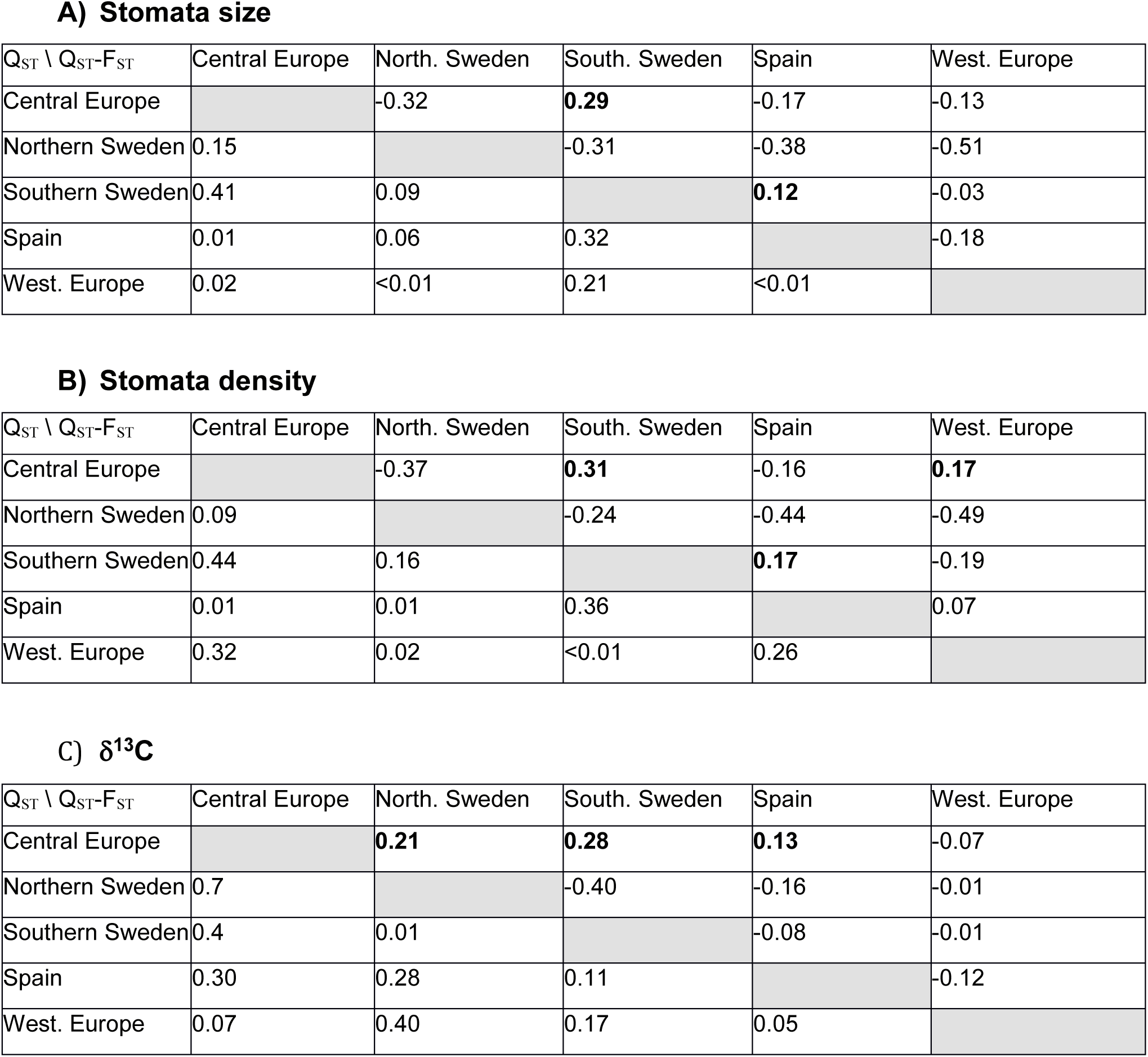
A-C: Patterns of regional differentiation depart from neutral expectations. Pairwise Q_ST_ estimates were derived from linear mixed models for all regions. Genome-wide, pairwise F_ST_ distribution was calculated based on 70,000 SNPs for all regions. In the top half of each table, the difference Q_ST_-F_ST_ for each pair of regions is shown. In the bottom half of each table the Q_ST_ estimate for each pair of regions is shown. Each table represents one phenotype as indicated by table headlines. Significant Q_ST_-F_ST_ differences are written in bold. The significance threshold is based on the 95th percentile of a distribution of maximum Q_ST_-F_ST_ values from 1000 random permutations of phenotypic data.

A) Stomata size
| $Q_{ST} \setminus Q_{ST}-F_{ST}$ | Central Europe | North. Sweden | South. Sweden | Spain | West. Europe |
| --- | --- | --- | --- | --- | --- |
| Central Europe |  | -0.32 | **0.29** | -0.17 | -0.13 |
| Northern Sweden | 0.15 |  | -0.31 | -0.38 | -0.51 |
| Southern Sweden | 0.41 | 0.09 |  | **0.12** | -0.03 |
| Spain | 0.01 | 0.06 | 0.32 |  | -0.18 |
| West. Europe | 0.02 | <0.01 | 0.21 | <0.01 |  |

B) Stomata density
| $Q_{ST} \setminus Q_{ST}-F_{ST}$ | Central Europe | North. Sweden | South. Sweden | Spain | West. Europe |
| --- | --- | --- | --- | --- | --- |
| Central Europe |  | -0.37 | **0.31** | -0.16 | **0.17** |
| Northern Sweden | 0.09 |  | -0.24 | -0.44 | -0.49 |
| Southern Sweden | 0.44 | 0.16 |  | **0.17** | -0.19 |
| Spain | 0.01 | 0.01 | 0.36 |  | 0.07 |
| West. Europe | 0.32 | 0.02 | <0.01 | 0.26 |  |

**Table 1 A-C: Patterns of regional differentiation depart from neutral expectations**
| $Q_{ST} \setminus Q_{ST}-F_{ST}$ | Central Europe | North. Sweden | South. Sweden | Spain | West. Europe |
| --- | --- | --- | --- | --- | --- |
| Central Europe |  | **0.21** | **0.28** | **0.13** | -0.07 |
| Northern Sweden | 0.7 |  | -0.40 | -0.16 | -0.01 |
| Southern Sweden | 0.4 | 0.01 |  | -0.08 | -0.01 |
| Spain | 0.30 | 0.28 | 0.11 |  | -0.12 |
| West. Europe | 0.07 | 0.40 | 0.17 | 0.05 |  |

Regional differences in climate may have imposed divergence in stomatal patterning. Thus, we estimated climatic distances between regions using estimates of regional effects extracted from a MANOVA. We did not observe significant correlations between adaptive phenotypic divergence (Q_ST_-F_ST_) and the climatic distance of the respective regions (Mantel test p>0.05 for each of the three traits). Regional divergence in δC^13^, stomata density and stomata size was therefore not proportional to climatic divergence.

## Discussion

### Genetic variation for stomata density and size segregates in A. thaliana

We used high-throughput confocal imaging to characterize stomata patterning in over 31,000 images from 870 samples collected from 330 genotypes. Our high-throughput pipeline could characterize stomata density and stomata size with a reliable accuracy, confirmed by high correlation with manual measurements. Broad-sense heritability and pseudo-heritability estimates for stomata density, which are 30% and 58%, respectively, are slightly lower than in a previous report of manually counted stomata diversity across a smaller sample chosen to maximize genetic diversity (Delgado et al., 2011). Despite the clear impact of environmental (random) variance on both observed phenotypes, stomata size and stomata density showed a strong negative correlation. This is consistent with earlier reports of studies manipulating regulators of stomata development (Doheny-Adams, Hunt, Franks, Beerling, & Gray, 2012; Franks et al., 2015), but also with studies analyzing stomatal trait variation in a wide range of species (Franks & Beerling, 2009; Hetherington & Woodward, 2003).

The extensive genomic resources available in *A. thaliana* enabled us to investigate the genetic basis of trait variation and co-variation, with the help of GWAS (Atwell et al., 2010). Much is known about the molecular pathways that control the differentiation of stomata in *Arabidopsis thaliana*, providing a set of candidate genes expected to control genetic variation in stomata patterning (Bergmann & Sack, 2007; Pillitteri & Torii, 2012). However, we did not detect any genomic region that associated with stomata density at a p-value beyond the Bonferroni-significance threshold. For stomata size, there was only one significant association on chromosome 4, albeit with very low minor allele frequency in a gene that has not been reported previously in stomata development. GWAS studies can detect small-effect loci only if they segregate at high frequency, whereas rare alleles only give detectable signals when they are of large effect (Korte & Farlow, 2013; Wood et al., 2014). Given that variance for both stomata size and stomata density is clearly heritable, the genetic variants controlling these traits are not causing strong association signals in GWAS. Theoretically, the presence of a large effect QTL impacting local adaptation can be masked by correction for population structure. However, not correcting for population structure is known to lead to a high number of false-positives and is thus not a reliable alternative (Vilhjálmsson & Nordborg, 2012). Nevertheless, we can conclude that variation in stomata patterning is controlled by a combination of i) alleles of moderate effect size segregating at frequencies too low to be detected by GWAS, and/or ii) alleles segregating at high frequency but with effect size too small to be detected and/or iii) rare alleles of small effect. In addition, it is possible that the effect of associated loci is weakened by epistatic interactions among loci. In *A. thaliana*, the genetic architecture of natural variation in stomata traits is therefore not caused by a handful of large effect variants but complex and polygenic.

Using MTMM analysis (Korte et al., 2012), we further investigated the impact of genetic variation on the negative co-variation between stomata size and density. This analysis revealed that genetic similarity does not influence the pattern of covariation. It implies that either multiple alleles act epistatically on the covariation, or that physical or environmental factors explain the correlation.

### Natural variation in stomata patterning can contribute to optimize physiological performance

Both stomata development and reactions to drought stress are being intensively investigated in *A. thaliana* (Bergmann & Sack, 2007; Krasensky & Jonak, 2012; Pillitteri & Torii, 2012; Verslues, Govinal Badiger, Ravi, & M. Nagaraj, 2013). Mutants in stomata density or size have recently been shown to have a clear impact carbon physiology (Franks et al., 2015; Hepworth, Doheny-Adams, Hunt, Cameron, & Gray, 2015; Hughes et al., 2017; S. S. Lawson, Pijut, & Michler, 2014; Masle, Gilmore, & Farquhar, 2005; Yoo et al., 2010; Yu et al., 2008). Yet, the relevance of natural variation in stomatal patterning for facing local limitations in water availability, had not been documented in this species so far. We provide here concomitant measures of morphological and physiological variation to examine the impact of variation in stomatal patterning on natural variation in WUE. By including genome-wide patterns of nucleotide diversity, our analysis presents two major findings: i) the decrease in stomata size associates with an increase in WUE in *A. thaliana* and ii) this pattern of co-variation has a genetic basis. This shows that, in *A. thaliana*, variation in stomata size has the potential to be involved in the optimization of physiological processes controlling the trade-off between growth and water loss. Interestingly, in the close relative *A. lyrata ssp. lyrata*, stomata were observed to grow smaller in experimental drought compared to well-watered conditions, which coincided with increased WUE (Paccard, Fruleux, & Willi, 2014). This suggest that the consequences of decreased stomata size are conserved in the genus.

While variation for stomata size and density is likely shaped by a complex genetic architecture that hindered QTL detection, we detected two regions in the genome that associated significantly with carbon isotope discrimination. Three previous QTL mapping analyses, including one between locally adapted lines from Sweden and Italy, identified 16 distinct QTLs controlling δ^13^C (Juenger et al., 2005; McKay et al., 2008; Mojica et al., 2016). One of these is caused by a rare allele in the root-expressed gene *MITOGEN ACTIVATED PROTEIN KINASE 12* (*MPK-12*), (Campitelli, Des Marais, & Juenger, 2016; Juenger et al., 2005). While QTL-mapping approaches can only reveal the variance shown by the parental lines, GWAS approaches fail to detect rare alleles unless they have a very strong impact. It is therefore not surprising that the loci that stand out in GWAS do not overlap with the QTL previously mapped. In fact, one of the mapping populations used the parental genotype Cvi-0, a genotypic and phenotypic outlier.

The two QTL we report here on chromosomes 2 and 4 add two novel loci, raising the number of genomic regions known to impact δ^13^C in *A. thaliana* to 18. The novel loci we report are locally frequent. Individuals carrying the minor alleles of both loci are almost exclusively from Southern Sweden and display significantly higher δ^13^C than other Southern Swedish accessions. However, we did not find any climatic factor significantly correlated with the allelic states of our QTLs. This suggests that other factors, like soil composition, play a role in drought adaptation. Alternatively, locally adapted alleles may not yet be fixed within the region.

Interestingly, the accessions with the minor allele associating with high δ^13^C in both QTL did not show decreased stomata size compared to other accessions. Multi-trait GWAS confirmed that these QTL are associated with δ^13^C variants that are independent of genetic variation for stomata patterning. We therefore can conclude that, stomata patterning is only one of the traits contributing to the optimization of WUE. A large array of molecular and physiological reactions is indeed known to contribute to tolerance to drought stress (Krasensky & Jonak, 2012; Verslues et al., 2013). The close vicinity of the chromosome 2 QTL to a non-synonymous mutation in a gene encoding an LEA protein, known to act as a chaperone when cells dehydrate, suggests one possible mechanism by which WUE might be optimized independently of stomata size and density (Candat et al., 2014; Eriksson, Kutzer, Procek, Gröbner, & Harryson, 2011; Reyes et al., 2005). Variation in rates of proline accumulation in the presence of drought stress or in nutrient acquisition in the root are also among the physiological mechanism that appear to have contributed to improve drought stress tolerance in this species (Campitelli et al., 2016; Kesari et al., 2012).

### Adaptive evolution of stomata patterning is suggested by the geographic distribution of genetic variation

Phenotypic variation for stomata patterning and carbon uptake is not uniformly distributed throughout the species range. All three phenotypes we report in this study were significantly differentiated between the five broad regions defined in our sample of 330 genotypes. We performed a comparison of phenotypic and nucleotide levels of divergence to evaluate the putative role of past selective events in shaping the distribution of diversity we report (Leinonen et al., 2013; Whitlock & Guillaume, 2009). Because these regions are not equally distant, F_ST_/Q_ST_ comparisons averaged over all populations may mask local patterns of adaptation (Leinonen et al., 2013). We therefore measured Q_ST_ between pairs of regions and compared them to the distribution of pairwise F_ST_, using permutations to establish the significance of outlier Q_ST_. This analysis showed that, for all three traits, differentiation between some regions was stronger than expected from genome-wide patterns of diversity, suggesting local adaptation. This is further supported by our finding that stomata density and stomata size correlated with climatic PCs, which are most strongly driven by temperature, humidity, solar radiation, and historic drought regimen.

The strongest Q_ST_-F_ST_ differences are found across regional pairs including Central Europe. Particularly, WUE is significantly differentiated between Central Europe and Spain as well as both Swedish regions, due to low WUE in Central Europe. It is tempting to speculate that the significantly lower WUE observed in Central Europe results from selection for life cycling at latitudes where two life cycles can be completed each year, as high WUE is usually associated with a reduction in photosynthetic rate (Blum, 2009; Field et al., 1983; Kimball et al., 2014). Interestingly, Central Europe and Southern Sweden are significantly differentiated for all three traits and Southern Sweden and Spain are significantly differentiated for both stomata traits. Combined with the fact that Swedish genotypes show the highest values for WUE, this suggests that stomata size is involved in drought adaptation of Swedish accessions. This result is somewhat counterintuitive because Sweden is not known to be a region experiencing intense drought. However, our result is supported by an independent study showing that Northern and Southern Swedish genotypes maintain photosynthetic activity under terminal drought stress longer than other, especially Central and Western European, accessions (Exposito-Alonso et al., 2017). Additionally, locally adapted genotypes from Northern Sweden (which showed high WUE in our study, as well) have been shown to display higher WUE than Italian genotypes (Mojica et al., 2016).

This regional difference in *A. thaliana* further coincides with the satellite measurements of historic drought regimen, which show that Sweden is a region where drought frequency is changing throughout the season: it is relatively more frequent in the early growing season (spring) than in the late growing season (summer). Drought episodes occurring earlier in the growth season may favor the evolution of drought avoidance traits (e.g. morphological or physiological stress adaptations) over that of escape strategies mediated by e.g. seed dormancy (Kooyers, 2015; Passioura, 1996). Indeed, in Northern Europe, increased negative co-variation between flowering time and seed dormancy suggested that the narrow growth season imposes a strong selection on life-history traits (Debieu et al., 2013). In Southern European regions, *A. thaliana* appeared to rely on escape strategies provided by increased seed dormancy (Kronholm et al., 2012). Taken together, this suggests that decreased stomata size and, consequently, increased δ^13^C have contributed to adaptation to water limitations in spring in a region where the narrow growth season leaves no room for escape strategies. Indeed, both stomata size and δ^13^C associate with historic drought regimen. For δ^13^C, however, this association disappears when genetic population structure is included as a covariate. This indicates that local adaptation for WUE might have contributed to shape current population structure.

Finally, the coarse regional contrasts used in the present study cannot resolve patterns of local adaptation occurring at a fine-grained scale within regions (as e.g. local adaptation to specific soil patches). In fact, we observe most variation for all three phenotypes within regions. It is therefore possible that we underestimate the magnitude of adaptive differentiation across the species’ European range, which could further explain why Q_ST_ / F_ST_ differences did not co-vary with environmental divergence in our dataset.

## Conclusion

This work provides a comprehensive description of the variation in stomata size and density that segregates throughout the European range of *A. thaliana*. It shows that stomata size covaries with water-use efficiency and may contribute to local adaptation. Several reports indicate that plants can also change stomatal development in water-limiting conditions (Fraser, Greenall, Carlyle, Turkington, & Friedman, 2009; Paccard et al., 2014; Xu & Zhou, 2008). Future work will have to investigate whether this variation in stomata size and number also contributes to adaptive plasticity to drought stress.

## Acknowledgements

We thank Swantje Prahl and Hildegard Schwitte for technical support in microscopy, Maria Graf for technical assistance in δ^13^C analysis, Anja Linstädter for advice in statistical analysis and Angela Hancock for critical comments on the manuscript. This research was supported by the Deutsche Forschungsgemeinschaft (DFG) through grant INST211/575-1 for the OPERA microscope, and grant ME2742/6-1 in the realm of SPP1529 “Adaptomics”, as well as by the European Research Council with Grant 648617 “AdaptoSCOPE”.

## Data accessibility

Raw image data and image analysis scripts are available upon request and will be stored in a Dryad repository upon acceptance. Phenotypic data is provided in the supplemental material and will be uploaded to the AraPheno database (https://arapheno.1001genomes.org, (Seren et al., 2017) and stored in a Dryad repository upon acceptance. Additionally, we provide an R Markdown file, which contains all figures (except GWAS and MTMM) and the corresponding R code used to create the figures and statistics in the supplemental material. GWAS scripts are available at https://github.com/arthurkorte/GWAS. MTMM scripts are available at https://github.com/Gregor-Mendel-Institute/mtmm. Genomic data used is publicly available in the 1001 genomes database (Alonso-Blanco et al., 2016)

## Author contributions

JdM, AK, and HD conceived the study. HD conducted the experiment and produced phenotypic data for stomata traits. TM and AW were responsible for δ^13^C measurements. GM provided data on historic drought regimen. JdM, AK and HD were responsible for the statistical analyses of the data. JdM and HD wrote the manuscript with significant contributions from AK, TM, AW and GM.

